# Spatial transcriptome of developmental mouse brain reveals temporal dynamics of gene expressions and heterogeneity of the claustrum

**DOI:** 10.1101/2023.04.12.536360

**Authors:** Yuichiro Hara, Takuma Kumamoto, Naoko Yoshizawa-Sugata, Kumiko Hirai, Song Xianghe, Hideya Kawaji, Chiaki Ohtaka-Maruyama

## Abstract

During the development of the mammalian cerebral cortex, numerous neurons are arranged in a six-layer structure with an inside-out fashion to form the neocortex and wire neural circuits. This process includes cell proliferation, differentiation, migration, and maturation, supported by precise genetic regulation. To understand this sequence of processes at the cellular and molecular levels, it is necessary to characterize the fundamental anatomical structures by gene expression. However, markers established in the adult brain sometimes behave differently in the fetal brain, actively changing during development. Spatial transcriptomes yield genome-wide gene expression profiles from each spot patterned on tissue sections, capturing RNA molecules from fresh-frozen sections and enabling sequencing analysis while preserving spatial information. However, a deeper understanding of this data requires computational estimation, including integration with single-cell transcriptome data and aggregation of spots on the single-cell cluster level. The application of such analysis to biomarker discovery has only begun recently, and its application to the developing fetal brain is largely unexplored. In this study, we performed a spatial transcriptome analysis of the developing mouse brain to investigate the spatiotemporal regulation of gene expression during development. Using these data, we conducted an integrated study with publicly available mouse data sets, the adult brain’s spatial transcriptome, and the fetal brain’s single-cell transcriptome. Our data-driven analysis identified novel molecular markers of the choroid plexus, piriform cortex, thalamus, and claustrum. In addition, we revealed that the internal structure of the embryonic claustrum is composed of heterogeneous cell populations.

## INTRODUCTION

The embryonic neocortex undergoes a dynamic formation process as cells differentiate and migrate to their intended locations ^1^. During the development of the brain, a large number of neurons assemble into detailed microstructures to form the neural circuits of the neocortex. Cellular processes such as cell division, differentiation, migration, and maturation are conducted by precise control of gene regulation. Gene expression provides a baseline for understanding such a series of dynamic cellular processes and the precise roles of individual microstructures at the cellular and molecular levels. Molecular markers have been reported for each of the anatomical structures. For example, *Rora* (RAR-related orphan receptor alpha) is specific to the thalamus ^2^, and *Tbr1* (T-box brain transcription factor 1) and *Cux2* (cut-like homeobox 2) are related to the deep and superficial layers of the neocortex, respectively ^3,4^. However, a precise molecular marker with a high level of specificity to the structure is not always available. Moreover, well-established markers in adults do not always maintain the same expression specificity in developing embryos. Overall, understanding the anatomical structure of the fetal brain at the molecular level remains a challenge.

Genome-wide expression profiling, such as RNA-seq ^5^ and CAGE ^6^, allows for the screening of marker candidates from a large pool of molecules. However, the preparation of target samples can be challenging when the size of the target area is microscopic. Single-cell RNA-sequencing ^7^ provides gene expression profiles, even from individual cells, but it requires the dissociation of cells, which leads to the loss of spatial information. Spatial transcriptome ^8^ is a recently developed approach for obtaining gene expression profiles from individual areas of a tissue section while maintaining spatial information. The spatial resolution depends on the technologies, from 2 to 100 μm ^9-11^, at the scale of dozens of cells to subcellular structure ^8^. A caveat in the interpretation of the spatial transcriptome is that spot-based gene expressions do not necessarily represent individual cells, as a single spot may contain multiple cells or extend across the border of adjacent cells. Understanding cellular expression requires computational estimation, including integration with single-cell transcriptome data and aggregation of spots at a cellular level.

Here, we performed a spatial transcriptome analysis of the developing mouse brain. We complemented our data with publicly available data sets, such as spatial transcriptome data of mouse adult brains and single-cell RNA-seq of fetal brains. Based on the spatiotemporally integrated transcriptome data that recapitulate the previously known anatomical structures, we found novel molecular markers of *Car12* (carbonic anhydrase 12), *Folr1* (folate receptor 1) for the choroid plexus, *Rprml* (reprimo-like) for the piriform cortex, *Hsd11b2* (hydroxysteroid 11-beta dehydrogenase 2) for the thalamus, and *Etl4* (enhancer trap locus 4) for the claustrum. Further integration of single-cell data sets and independent experiments uncovered an internal substructure of the embryonic claustrum consisting of heterogenous cell types.

## RESULTS

### Spatial gene expression landscape of the early mouse brain

To understand gene expressions in individual anatomical areas of the developing brain, we performed spatial gene expression profiling using the Visium platform (10x genomics). We obtained coronal sections of the forebrain in two stages (E17 and P0) of developing Institute of Cancer Research (ICR) mice for analysis. The resulting data was complemented by publicly available data of an adult C57BL/6 mouse brain produced by the same platform. In the E17 and P0 profiles, 7,560 and 6,957 genes were detected in the median, respectively, whereas 6,048 genes were found in the adult profile. We confirmed that the expressions of Satb2 (special AT-rich sequence binding protein 2), *Slc6a4* (solute carrier family 6, member 4), *Gbx2* (gastrulation brain homeobox 2), and *Neurod1* (neurogenic differentiation 1) genes, known markers of excitatory neurons of the neocortex upper layers, ventral posterior (VP) complex of the thalamus, non-VP part of the thalamus, and hippocampus, respectively, were consistent with previous reports^12-16^ and in situ hybridization (ISH) results (Figs. S1, S3, S4, S5). These results confirmed that our spatial gene expression profiles are reasonably aligned with the current knowledge.

Next, we integrated the expression data of the individual spots from the three stages to obtain spatiotemporally coherent gene expression profiles by employing the anchor-based approach, implemented in Seurat ^17^. The gene expressions of individual spots were subjected to clustering and visualization by dimension reduction with Uniform Manifold Approximation and Projection (UMAP) (Fig. 1a, see Methods). In all stages of E17, P0, and adults, clusters c1, c23, c14, and c15 of the resulting 25 spatial spot clusters corresponded to regions expressing *Satb2* as the neocortex upper layers (II/III), *Slc6a4* as the VP, *Gbx2* as the non-VP of the thalamus, and *Neurod1* as the hippocampus, respectively (Figs. 1b, S1– S5). We found that the spot clusters agreed well with the overall anatomical structure.

**Fig. 1.**
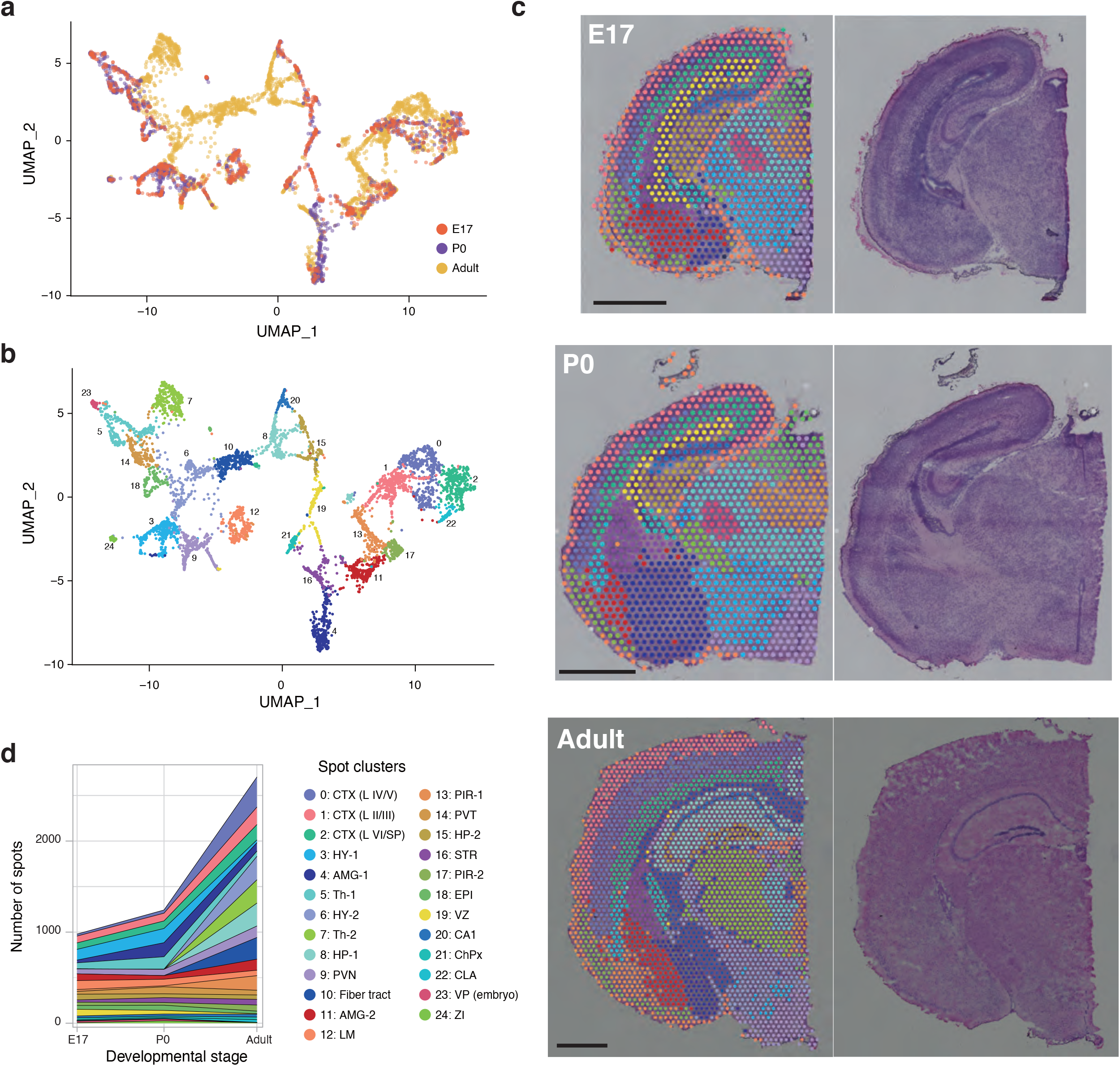
Overview of the spatial transcriptome analysis of the developing mouse brains. **(a**,**b)** UMAP projection of the spots based on the integrated data, where spots were annotated by sample stages (a) and clusters (b). **(c)** H&E staining of tissue sections overlayed with the spots. **(d)** Number of spots in the individual clusters that were produced by the integration analysis.

Then, we manually annotated the clusters based on their locations in the tissue sections and expressions of known marker genes (Figs. 1c, S6). The majority of the clusters appeared in all three stages, with some exceptions, where the number of spots corresponding to the clusters reflect the physical sizes and changes in anatomical structure (Fig. 1d). The adult-specific clusters c6, c7, c8, c10, and c13, correspond to HY-2, Th-2, HP-1, Fiber tract, and PIR-1, respectively. These clusters are featured by the presence of genes expressed in matured neural circuitry such as synaptic function-related genes, neurotransmitters, and myelin sheath component genes. Notably, Tnnt1 and Prkcd are specifically expressed in c7 but absent in the embryonic stages. In contrast, c19 and c23 are unique to developing stages and correspond to the ventricular zone (VZ) and VP complex of the thalamus, respectively. The VZ is known to contribute to neuron production by localizing neural progenitors during embryonic development and is lost in adults ^18^. These observations underline substantial changes in the gene expressions in these areas, and it is reasonable to consider these clusters as specific to the developmental stages. Besides the five and two clusters only observed in the adult and the developing stages, the remaining clusters agreed across all three integrated stages.

### Data-driven exploration of molecular marker candidates

Next, we explored novel marker candidates in a data-driven manner. We performed differential expression analysis between the focused clusters and the rest, for the E17 and P0 stages. The genes with significantly high expression in the specific clusters contained novel marker candidates, as well as some known markers.

The expression profiles of the identified genes are visualized as a heatmap with known marker genes (Fig. 2). The heatmap shows that the expressions between E17 and P0 are largely consistent. For example, the *Slc6a4* and other VP markers, *Rasd1* and *Zmat4*, were substantially active in c23 (VP), in both stages. Their weak presence in c5 (non-VP part of the thalamus) was also commonly observed in both stages. We also found that Nexn (nexilin), *Wnt9b* (wingless-type MMTV integration site family, member 9B), and *Gckr* (glucokinase regulatory protein) show a similar expression specificity to VP, indicating their potential utility as VP markers. We chose some of the novel marker candidates (indicated by the red bold font in Fig. 2) in the developing stages and examined them individually.

**Fig. 2.**
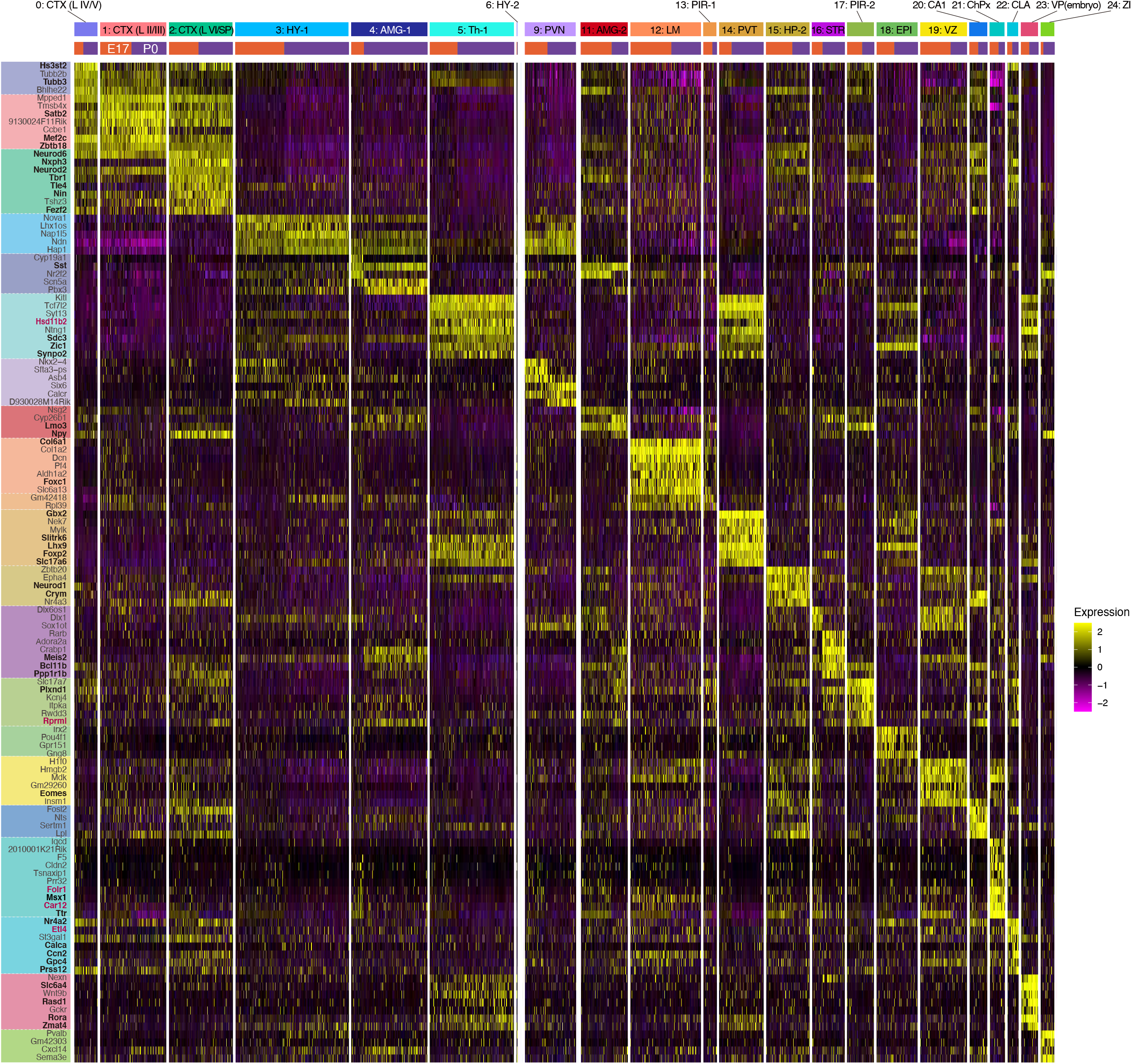
Heatmap of the genes specifically expressed in the individual clusters. Each row and column represent the gene symbol and spot, respectively. The spots were classified by the clusters to which they belong (upper bars; see Fig. 1d) and experiments from which they were produced (lower bars). Gene symbols in black bold and red bold represent molecular markers reported previously and novel markers discovered in this study, respectively.

### Novel molecular markers in the choroid plexus, thalamus, and piriform cortex

The choroid plexus (corresponding to c21 of our spot clusters) is the epithelial tissue within the ventricles of the brain that arises from the cortical hem, and it produces cerebrospinal fluid (CSF), which plays an important role in the development and function of the adult brain. Msx1, a known choroid plexus marker, is highly active in c21, but it is also expressed in leptomeninges (c12) at E17 and P0 (Fig. 2). The choroid plexus initially forms when the pia matter invaded the ventricles and adhered to the layer of ventricular ependymal cells, and it shares a developmental origin with the leptomeninges. Therefore, it is reasonable that Msx1, which is highly expressed in the choroidal choroid, is also expressed in a cluster of the leptomeninges (c12).

Our analysis identified two genes, *Folr1* and *Car12*, as novel molecular marker candidates of the choroid plexus, and their expression in leptomeninges is less frequent (Fig. 2). To examine their expression in an independent experiment, we performed ISH across the developmental stages, from E11 to P0 (Fig. 3A). We found their exclusive presence in the choroid plexus with an absence in leptomeninges during the stages of E17 and P0, which confirmed our observation from the spatial transcriptome data and underline their utilities as biomarkers. Notably, *Folr1* was shown after the stage of E11, and *Car12* was shown after E14.

**Fig. 3.**
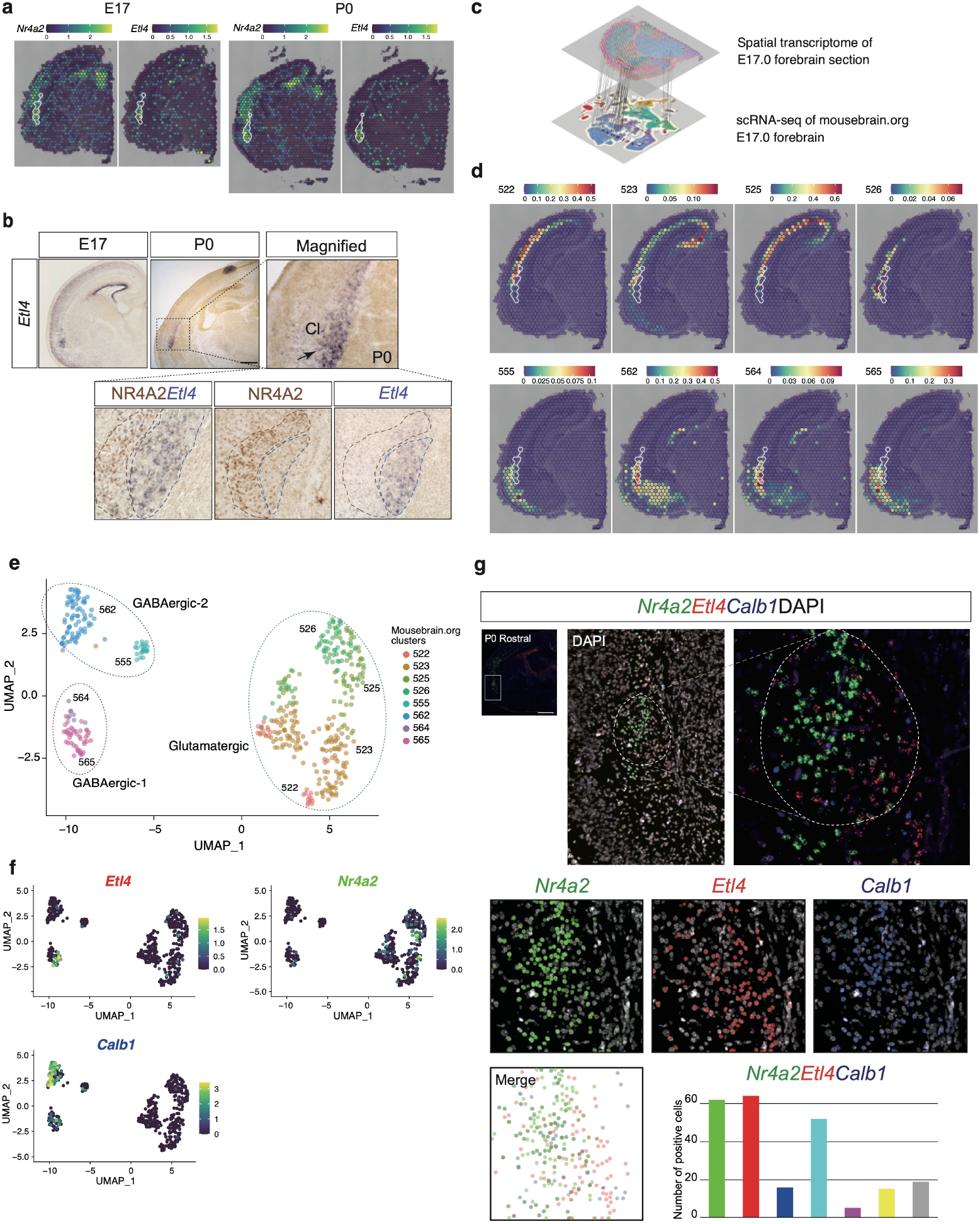
Novel markers of the choroid plexus, thalamus, and piriform cortex. (a) *Folr1* and Car12 were identified as novel markers of the choroid plexus in c21. Gene expression patterns at P0 cortex by Visium (left). In situ hybridization was performed to examine *Folr1* and Car12 expression at various stages of development from E11 to P0, confirming their expression in the choroid plexus (2^nd^ and the following columns). (b) *Hsd11b2* and *Rprml* were identified as novel marker genes for the thalamus (c5) and piriform cortex (c17), respectively. Gene expression patterns at P0 cortex by Visium (left). In situ hybridization was performed to examine *Hsd11b2* and Rprlm expression at various stages of development from E11 to P0, confirming their expression in the choroid plexus (2^nd^ and the following columns). ChPx: choroid plexus, VP: ventral posterior nucleus, Th: thalamus, SP: subplate, HP: hippocampus, PIR: piriform cortex, AMG: amygdala.

The thalamus (c5 and c23), which first projects axons to the cortex and relays sensory information, such as visual, auditory, and somatosensory information, to the cortex, is anatomically composed of numerous neural nuclei. During the development of mesencephalon, the thalamus is first regionalized by the expression of various transcription factors and is further patterned by concentration gradients of signaling molecules such as *Shh* (sonic hedgehog), *Wnt*, and *Fgf8* (fibroblast growth factor 8) ^19^. The thalamus has a relay nucleus that transmits somatosensory information to the cortex (called the VP nucleus; c23), and a thalamus markers, *Rora* and *Slc6a4* are known to be highly expressed in the VP within the thalamus. In contrast, we discovered that the gene *Hsd11b2* is expressed at a higher level in the non-VP part of the thalamus (c5) than in the VP part (c23) (Fig. 2). Our ISH experiments confirmed its unique presence in the non-VP part of the thalamus and its absence in the VP part (Fig. 3b). The contrastive expression of *Hsd11b2* against *Rora* or *Slc6a4*(Fig.S1) indicates the utility of their combination in delineating the internal structure of the thalamus.

The piriform cortex (PIR-2, c17) is one of the structurally simplest and evolutionarily oldest cortices, called the paleocortex. After the olfactory epithelium and olfactory bulb, it constitutes the third stage of the odor information processing pathway. In contrast to the neocortex consisting of six layers, the piriform cortex has a three-layered structure, where afferents and efferents are established in a layer-specific manner ^20^. Diodato et al. have identified layer-specific genes in the piriform cortex by laser-capture micro-dissection and RNA deep sequencing ^21^. They found genes enriched in the three layers but did not examine their expression in the embryonic stage. Our analysis revealed enriched genes in the piriform in embryonic stages, including Slc17a7, Plxnd1, Itpka, Rwdd3, *Rprml*, Lmo 3, and Btbd 3. Lmo3 and Btbd3 were among the genes that Diodato et al. reported to be expressed in the piriform in the adult stage, and Plxnd1 has been reported to be expressed in the embryonic piriform ^22^. However, the other genes were newly identified as piriform markers of the embryonic stage in this analysis. We found that *Rprml i*s expressed in the piriform cortex (c17), hippocampus (c20), and subplate layers (c2), similarly to Plxnd1 (Fig. 3c). Moreover, Itpka showed greater specificity only under the developmental stages (Fig. S10).

### Etl4, a highly specific maker of the claustrum, delineates its inner structures

The claustrum is a thin sheet-shaped region in the basolateral forebrain that is thought to be involved in four main functions: sleep regulation, consciousness, attention and saliency, and memory ^23-25^. This is supported by the fact that the prefrontal cortex is in close contact with the claustrum ^26^. According to BrdU-based birth date analysis of the cortical progenitors, claustrum neurons are born around embryonic day 12.5, which is later than adjacent subplates and endopiriform nuclei ^27^. Most cortical areas have projections to the claustrum. There are strong projections from the prefrontal cortex, including the orbitofrontal cortex (OFC), prelimbic cortex, anterior cingulate cortex (ACC), and second motor cortex, and from the insula and temporal cortex to the claustrum, as well as output from the claustrum to the majority of the prefrontal cortex ^26^. However, owing to the absence of clear structural features and the variety of its shapes among mammals ^28^, it has been challenging to define precise anatomical boundaries. The definition of the claustrum is relatively broad and consists of distinct subregions such as the dorsal claustrum, ventral claustrum, and the dorsal endopiriform nucleus (DEn) ^29^. As in other cortical areas, DEn is approximately 90% glutamatergic neurons and expresses *Nr4a2* (nuclear receptor subfamily 4, group A, member 2; also known as Nurr1), a well-known marker. Notably, most of the identified genetic markers are expressed in the preoptic region and other areas, such as the DEn and subplate, and in deep layers of the lateral neocortex in rodents ^30-33^. Our data support this expression pattern, as *Nr4a2* was found in the subplate neurons (c2) and VZ (c19), indicating the limitation of *Nr4a2* as a claustrum/DEn marker.

The limited knowledge of the anatomical features of the claustrum has hindered the precise elucidation of its substructures. Our spatial transcriptome data identified the *Etl4* gene as a potential molecular marker whose expression is more specific for claustrum/DEn than that of *Nr4a2* at both stages of E17 and P0 (Figs. 2, 4a). We examined their expressions by performing ISH for *Etl4* and immunohistochemistry for NR4A2. Notably, *Etl4* and NR4A2 were found in distinct areas within the claustrum/DEn (Fig. 4b). The *Etl4* signal marked the inner area, while NR4A2 was found in the outer area, with clear contrast. *Gng2* and *Ntng2* are also known as claustrum molecular markers ^34,35^ but did not show specific expression in it, in the developing brain. This result demonstrates that *Etl4* indicates a distinct part of the claustrum that cannot be illustrated by the known markers and the presence of a substructure of the claustrum that has not been reported previously in the developing brain.

**Fig. 4.**
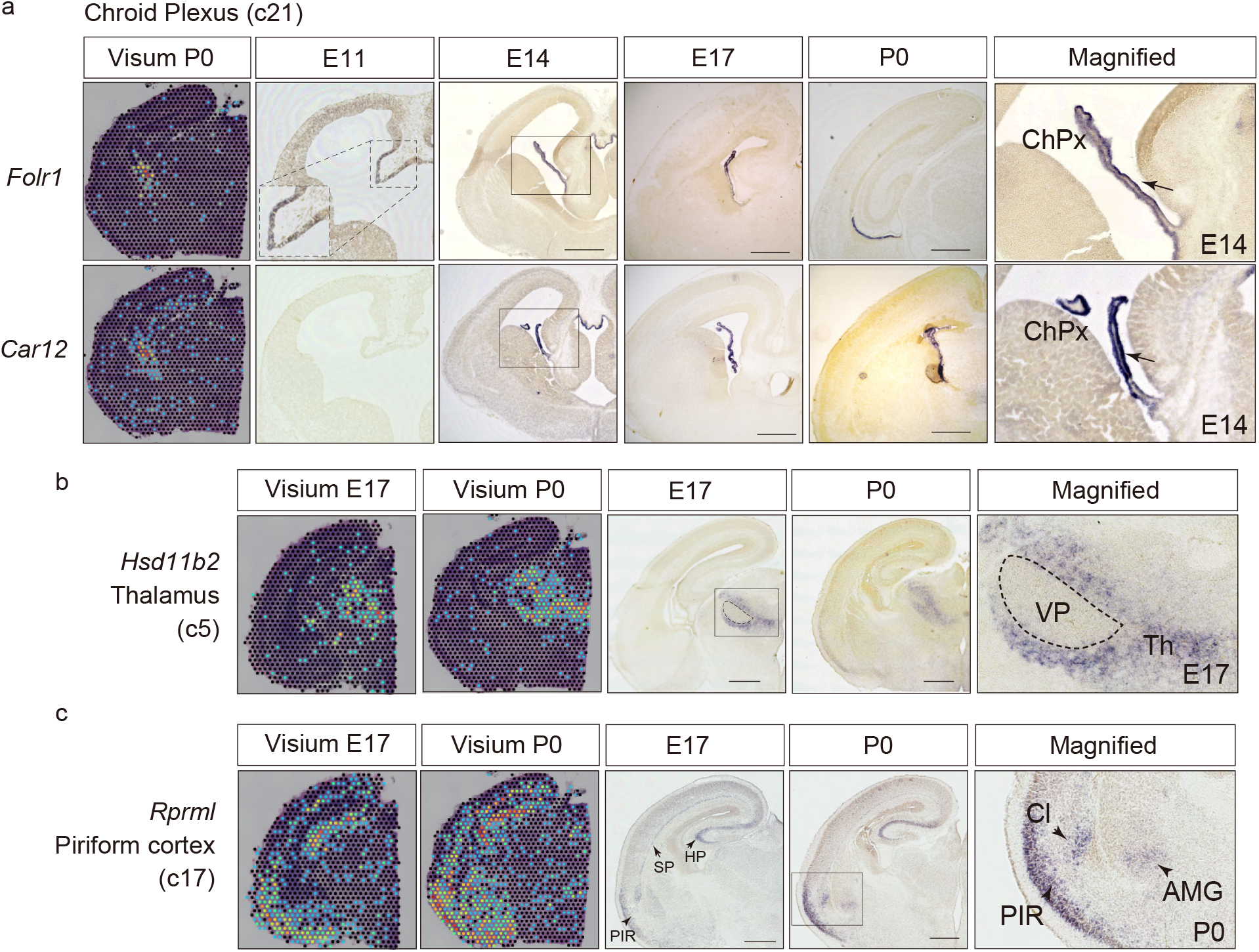
Cell populations in claustrum revealed by integration analysis of spatial transcriptome and scRNA-seq. **(a)** Expression of a known marker *Nr4a2* (Nurr1) and a novel marker *Etl4* in the claustrum of developing mice demonstrated by the spatial transcriptome analysis. The area with a white outline indicates the claustrum. **(b)** Expressions of NR4A2 (immunohistochemical staining) and *Etl4* (in situ hybridization) within the claustrum. Cells in the left area express NR4A2 but not *Etl4*, and those in the right area primarily express *Etl4*. Cl: claustrum **(c)** Schematic view of anchor-based integration between our spatial transcriptome data and the scRNA-seq data from mousebrain.org. **(d)** Visium spots of the E17 slice labeled by prediction scores that measured the confidence level of the cell types referring to the scRNA-seq cell clusters from mousebrain.org. Each slide contains the projection scores of a cell cluster that were positive in the claustrum area (Visium cluster 22). The same area indicated by the white line in (a) is shown. **(e)** UMAP projection of single-cell gene expression profiles of the cell clusters shown in (d). The cells were classified into three groups in the UMAP projection: one glutamatergic neuron group and two GABAergic neuron groups. **(f)** Known and novel markers of the cells associated with claustrum. **(g)** Multiplexed fluorescence in situ hybridization of the claustrum/DEn at P0 cortex. Top: max projection images labeled with *Nr4a2* (C1), *Etl4* (C2), and *Calb1*(C3) channels. Overview of hemisphere (left), expansion of the white square (middle), and expansion of the claustrum/DEn region (right). Middle: Each signal (plot>1)-positive cells plotted in spatial coordinates. Three channel-mixed plot show in right. Bottom: Quantification of fluorescent-positive cells within the claustrum/DEn region.

### Distinct GABAergic inhibitory neurons comprising the claustrum/DEn inner structure

To understand the cell populations within the claustrum/DEn in the developing brain, we used publicly available single-cell RNA-seq data, an atlas of the developing mouse brain in mousebrain.org ^36^, and integrated the data into our spatial transcriptome data. We retrieved the expression data of the single cells with their cluster information and projected them onto the spatially arranged spots of the E17 (Fig. 4c) by employing an anchor-based approach ^37^. We found eight clusters of the single cells from the atlas that match the claustrum/DEn (Fig. 4d). Notably, the eight cell clusters were classified into three major groups in a UMAP plot based on their gene expression profiles: one group representing excitatory glutamatergic neurons and the other two representing inhibitory GABAergic ones. Two of the three clusters were characterized by *Etl4* and *Nr4a2* expressions, where *Etl4* positive cells primarily belong to one of the GABAergic neuron groups, GABAergic-1 (Fig. 4e,f). This indicates that GABAergic and glutamatergic neurons reside within the inner and out regions, respectively. We performed immunostaining of NR4A2 against GABAergic neuron labeled mice by TdTomato and confirmed the preferences of glutamatergic and *Etl4* positive neurons to the outer and inner structures, respectively (Fig. S8).

A remaining question is the location of the other GABAergic neurons (GABAergic-2) and if they reside within the inner or outer regions. We found that *Calb1* (calbindin 1) was a highly expressed gene in GABAergic-2 compared with the other two (Fig. 4f). We performed a fluorescence ISH (FISH) assay with single molecule sensitivity, RNAscope (Advanced Cell Diagnostics), and found that the inhibitory GABAergic-2 neurons reside in the claustrum in both the inner and outer regions (Fig. 4g).

## DISCUSSION

We performed spatial transcriptome analysis to explore molecular features of anatomical microstructures in the developing mouse brain. We found that our gene-expression-based clusters recapitulate the current knowledge of anatomical structures. The data-driven exploration of molecular marker candidates and subsequent validation experiments led to the identification of *Car12* and *Folr1* as novel molecular markers of the choroid plexus, *Hsd11b2* as of the thalamus, *Rprlm* as of the piriform cortex, and *Etl4* as of the claustrum/DEn. While adult brains have been studied with the establishment of spatial transcriptome technology ^11^, our study focusing on the developing brain led to these novel findings. Furthermore, an extensive computational analysis using the single-cell gene expression atlas ^36^ led to the discovery of heterogenous neuronal cell types, a specific subtype of GABAergic neurons comprising a substructure of the claustrum.

The marker genes identified in this study exhibit unique specificities of spatiotemporal gene expressions, and they may be used solely or complementarily with the known markers. The choroid plexus is the epithelial tissue within the ventricles of the brain, which plays an important role in the development and function of the adult brain by producing CSF. WNTs and BMPs are known to be expressed in the telencephalon ^38,39^, and a recent study reported that precise regulation of Wnt signaling is critical for normal choroid plexus development ^40^. We found that *Msx1*, a known choroid plexus marker in adults, is also specific to the choroid plexus in the developing brain, but it is also expressed in the leptomeninges. The novel choroid plexus markers, *Folr1* and *Car12*, were absent in leptomeninges, which indicates their strong specificity to the choroid plexus. Notably, *Folr1* was expressed at the developing stages after E11, whereas *Car12* was expressed after E14. *Folr1* is applicable in broad stages of development, before and after the choroid plexus maturation.

The thalamus consists of numerous neuronal nuclei, and the VP is its substructure, corresponding to a relay nucleus that transmits somatosensory information to the cortex, which is characterized by the expression of the *Rora* gene. *Tcf7l2* and *Gbx2* are known markers of non-VP regions in the embryonic thalamus ^14,41,42^, but they have limited specificity as found in the paraventricular nucleus (c14, Fig. 2). The novel marker of the non-VP thalamus, Hsd11b2, exhibited greater specificity. Moreover, it becomes fully silent in adults, whereas *Tcf7l2* and *Gbx2* remain active in adults (Fig. S9). In both anatomical and developmental aspects, *Hsd11b2* has greater specificity than the known ones.

The piriform cortex constitutes the third stage of the odor information processing pathway, after the olfactory epithelium and olfactory bulb, and it structurally belongs to the paleocortex, one of the structurally simplest and evolutionarily oldest cortices. Although Plxnd1 was previously reported as its marker gene, we found that *Rprml* shows a compatible expression specificity, with some expression in the hippocampus (c20) and subplate layers (c2), like Plxnd1. Although subplate neurons are located in the lowest layers of the neocortex (6b), the sharing of these molecules may imply a common feature of these areas, for example, a developmentally similar origin to the paleocortex. We also found that Itpka only has a greater specificity under development (Fig. S10).

The claustrum is known for the elongated subcortical area, which receives input from other regions, with interconnection to many areas within the cortex (Madden et al. 2022). The adult claustrum has a close connection with the ACC in rodents ^43-45^ and controls specific behavioral and cognitive functions with input from the ACC ^46,47^. Several functions involved in top-down cognitive processing have been proposed as its roles ^48-50^. We found a novel marker for the claustrum, *Etl4*, which is specific to inhibitory neurons and demonstrates greater specificity of expression patterns than the known marker *Nr4a2*, specific to glutamatergic neurons (Fig. 4a,b,e,f). The co-expression pattern of these two markers may help clarify the unambiguous shape of the claustrum (Fig. 4b). The relevance of *Etl4* in brain development and its molecular functions remain to be explored. It is expressed in a wide range of tissues but at a low level, and its human homolog, KIAA1217, has been poorly characterized. A recent study reported its genetic association with vertebral malformation ^51^, which indicates its contribution to development.

The identification of the novel marker for claustrum extends to the investigation of the substructure of the area in the developing brain. The integration of our spatial transcriptome data with the public scRNA-seq data led to a hypothesis of heterogenous cell populations within the claustrum, in particular, three clusters of cells: one *Nr4a2*-positive glutamatergic neuron cluster and *Etl4*- and *Calb1*-positive inhibitory neuron clusters (Fig. 4e,f). The idea of cellular heterogeneity is aligned with the presence of a unique migration mode of some neurons forming the claustrum ^52^. Our immunostaining and ISH experiments further uncovered the inner/outer structure of these three clusters that were depicted by inhibitory/excitatory neurons within the claustrum (Fig. 4b,g). This substructure may be worth comparing with those in adult brains that have been observed in several studies to elucidate the structural development of the claustrum. An electrophysiological study showed five subtypes of neurons, two spiny projection neurons and three aspiny interneurons ^46^. They found that the input from the ACC to the claustrum, composed of complex microcircuitry, encodes an anticipatory top-down signal required for optimal performance accuracy ^47^. At the molecular level, a study using scRNA-seq and mFISH showed that the claustrum in adult mice comprises two excitatory glutamatergic neuron subtypes that are differentiable from the surrounding cortex, forming a core-shell structure ^53^. This core-shell structure of the excitatory glutamatergic neurons might be similar to the inner/outer structure in developing mice, where the latter contains inhibitory neurons. Our result uncovered a heterogenous cell population of the claustrum in the stage of E17, but the relationship between its internal structures during and after development remains to be elucidated.

We explored the developing brains in mice by spatial transcriptome analysis and its integration with single-cell expression data, which uncovered novel markers of four anatomical regions as well as a delineated substructure of the claustrum. The results expand our knowledge of gene expressions underpinning brain development and provide insight into neuroanatomy based on gene expression dynamics.

## Acknowledgments

Computations were partially performed using the NIG supercomputer at ROIS National Institute of Genetics. H.K. and C.O.-M. were supported by the Takeda Science Foundation, and T.K. was supported by a Leading Initiative for Excellent Young Researchers (LEADER), grant number 2020L0019. This work was also supported in part by the JSPS KAKENHI-Grants (17K07428, 19H04795, 20H03270 to C.O.-M.), and AMED under Grant Number JP21gm1310012, Y2018 Research Grant from the Takeda Science Foundation, the Naito Foundation, FY2020 Research grant from the Novartis Foundation, the Brain Science Foundation, FY2021 Research Grant from the Yamada Science Foundation, the KOSE Cosmetology Research Foundation, the Mitsubishi Foundation, and the Astellas Foundation for Research on Metabolic Disorders to C.O.-M.

## Methods

### Spatial gene expression profiling

All animals were treated in accordance with the Tokyo Metropolitan Institute of Medical Science Animals Care and Use Committee guidelines. Pregnant ICR mice were purchased from Japan SLC Inc. (Hamamatsu, Japan). The brains of the embryos were harvested at E17.5 and P0 and directly embedded using an OCT compound. Embedded brains were stored at −80°C until sectioning. Frozen sections were prepared from each brain using thicknesses of 10 μm, following the manufacturer’s protocol. The sections were placed on the Visium Spatial Gene Expression Slide (10x Genomics, USA) and processed with the Visium Spatial Gene Expression Reagent kit (10x Genomics, USA) according to the manufacturer’s instructions. H&E brightfield and FL staining images were collected as mosaics using a Keyence BX-800 microscope (Keyence, Japan). The 10x Visium array covered about 25% (E17) and 30% (P0) of the spots. The resultant libraries were sequenced at Macrogen Inc. using the HiSeqX platform (Illumina, USA) with 150bp paired-end sequencing.

### Data processing of spatial gene expression

FASTQ files were processed using Space Ranger 1.2.0 (10x Genomics Inc., USA) with default parameter settings referring to the gene annotations of the mouse mm10 genome assembly provided by 10X Genomics (refdata-gex-mm10-2020-A), resulting in production of the matrixes of gene expression profiles of the individual cells. Visium spatial transcriptome data of a coronal section of an adult mouse cerebrum that was publicly available were integrated into our analyses. Its FASTQ files were retrieved from the 10X Genomics website (https://www.10xgenomics.com/resources/datasets). The gene expression profiles of the E17, P0, and adult samples were processed using the Seurat v3.2.3 platform ^36^. Read count matrixes were normalized before dataset integration by employing the anchor-based approach ^36^ with the Seurat FindIntegrationAnchors and IntegrateData functions. The spots of the three samples were grouped based on their expression profiles by employing the Shared Nearest Neighbor approach using the Seurat FindNeighbors and FindClusters functions setting at a resolution value of 0.95, resulting in 25 clusters. UMAP was performed for dimension reduction of the gene expression profiles of the individual spots using the Seurat RunUMAP function. 2D UMAP plots were visualized using the Seurat DimPlot function and the Loupe Browser application. Marker genes, those with significantly higher expression levels in a cluster than the others, were detected using the Seurat FindMarkers. The clusters were manually annotated referring to the ‘known’ marker genes that were included in those that were automatically inferred.

### Integration with single-cell RNA-seq

The scRNA-seq expression values of a developing mouse brain were retrieved from mousebrain.org ^36^ (http://mousebrain.org/development/downloads.html). The dataset, which was stored in the loom format, was converted into the ReadVelocity function implemented in the SeuratWrapper R library using the Seurat as. Seurat function. We extracted the expression profiles of forebrain neural cells at stage E17 from the dataset to integrate them into the spatial expression profiles of the E17 forebrain slice. The cluster annotation of single cells was used for the following analysis.

The expression values of the spots of the E17 slice were normalized by employing the SCTransformation approach ^54^ using the Seurat SCTransform function. These spots were then clustered based on gene expression profiles using the Seurat FindNeighbors and FindClusters functions, and dimension reduction of the gene expression profiles was performed using the Seurat RunPCA and RunUMAP functions. The expression values of E17 forebrain single cells were processed in the same way. Integration of the scRNA-seq dataset into spatial transcriptome one was performed using the Seurat FindTransferAnchors function ^36^, where an anchor-based approach was performed by querying the spatial gene expression profiles and referring to single-cell gene expression profiles. The integrated dataset was further processed using the Seurat TransferData function, which yielded a prediction score (a confidential index of cell type prediction) for the individual single-cell clusters at every spot. We chose the single-cell clusters that exhibited high prediction scores in the spots of Visium clusters 21 (choroid plexus) and 22 (claustrum). The prediction scores were visualized by overlaying them onto the Visium spot using the Seurat SpatialFeaturePlot function (Figs. 4d, S2a).

The selected single-cell clusters were visualized using UMAP plots with the aforementioned procedures, and the clusters were grouped by visual inspection based on the UMAP plot. Among these groups, we searched for marker genes that were more highly expressed in one group than the others, using the Seurat FindMarkers function.

### In situ hybridization, fluorescence in situ hybridization, and immunostaining

ISH was carried out using digoxigenin-labeled riboprobes as described previously (21). Coronal sections of wild-type brains (E14–P0) were used for this experiment. For the detection of *Nr4a2, Etl4*, and Kcnip1 mRNA in Fig. 4g, we used the RNAscope™ Multiplex Fluorescent V2 kit (Advanced Cell Diagnostics) and Fluorescein, Cy3, and Cy5 TSA fluorophore (Akoya Biosciences) according to the manufacturer’s protocol with minor modifications. Briefly, cortical sections (20 μm) were obtained using a cryostat. The slides were subsequently stored at –80°C until use. Coronal brain sections were baked at 60°C for 30 min, fixed with 4% paraformaldehyde in PBS on ice, washed with 100% ethanol, then treated with hydrogen peroxide for 10 min at room temperature. After washing with water, the sections were boiled for 5 min in a target retrieval buffer (Advanced Cell Diagnostics). After target retrieval, sections were washed with water followed by 100% ethanol, and then they were left to dry overnight. Next, the sections were treated with the protease for 10 min at 40°C and exposed to the *Nr4a2, Etl4*, and Kcnip1 probes (Advanced Cell Diagnostics) for 2h at 40°C before signal amplification. The fluorescence signal was further amplified using the TSA plus system (Akoya Biosciences) before detection. Sections were finally washed and treated with DAPI for 2 min before mounting in Fluoro-KEEPER Antifade Reagent (Nakalai Tesque).

For immunostaining, the embryonic brains were dissected and fixed in 4% paraformaldehyde/PBS overnight at 4°C. The tissues were cryoprotected in 15% sucrose/PBS overnight, followed by 30% sucrose/PBS overnight at 4°C. The brains were then embedded in an OCT compound (Tissue Tek) and cut into 20 μm-thick sections using a HM550 microm (PHC). The sections were soaked in PBS for 5 min, and pre-incubated with 0.3% TritonX-100/PBS for 15 min, which were then incubated overnight at 4°C with primary antibodies, anti-Nurr1(1:100) (AF2156, R&D systems), diluted with PBS containing 0.5% skim milk. After washing three times with PBS, the sections were incubated with species-specific anti-IgG antibodies conjugated to AlexaFluor 488. Then, sections were mounted with Fluoro-KEEPER Antifade Reagent (Nakalai Tesque) after DAPI staining (5 μg/ml, Sigma-Aldrich). Images were captured using a Stellaris 5 (Leica) or FV3000 (Evident) confocal microscope.

## Data availability

Sequence reads were deposited in the NCBI Short Read Archive under BioProject accession number PRJNA949444,, and the processed data with SpaceRanger and slide images were deposited in the NCBI Gene Expression Omnibus under the accession number GSE229508.

## Figure legends

Figure legends (for supplemental Figures)

Fig. S1. Expression pattern of known marker genes

Top: Cluster (top) and gene expression (middle) patterns at E17 cortex by Visium. Bottom: Expression of Satb2 (https://developingmouse.brain-map.org/experiment/show/100091731), Slc6a4 (unknown age, between E18 and P4: https://developingmouse.brain-map.org/gene/show/15342), *Gbx2* (https://developingmouse.brain-map.org/experiment/show/100091700), and NeuroD1 (https://developingmouse.brain-map.org/experiment/show/100091691) in E17 brain from Allen Mouse Brain Atlas.

Fig. S2. Temporal expression pattern of Satb2

Top: Cluster (top/middle) and gene expression (bottom) patterns of Satb2 from E17 to the adult cortex by Visium. Bottom: Expression of Satb2 from Allen Mouse Brain Atlas. E17 (https://developingmouse.brain-map.org/experiment/show/100091731),P1 (https://developingmouse.brain-map.org/experiment/show/100092087), and P28 (https://developingmouse.brain-map.org/experiment/show/100093561).

Fig. S3. Temporal expression pattern of Slc6a4

Top: Cluster (top) and gene expression (middle) patterns of Slc6a4 at E17 and P0 cortex by Visium. Bottom: Expression of Slc6a4 from Allen Mouse Brain Atlas. (E17, unknown age, between E18 and P4) https://developingmouse.brain-map.org/gene/show/15342

Fig. S4. Temporal expression pattern of *Gbx2*

Top: Cluster (top) and gene expression (middle) patterns of *Gbx2* from E17 to adult cortex by Visium. Bottom: Expression of *Gbx2* from Allen Mouse Brain Atlas. E17 (https://developingmouse.brain-map.org/experiment/show/100091700), P1 (https://developingmouse.brain-map.org/experiment/show/100091990), P28 (https://developingmouse.brain-map.org/experiment/show/100093579).

Fig. S5. Temporal expression pattern of neuroD1

Top: Cluster (top) and gene expression (middle) patterns of NeuroD1 from E17 to adult cortex by Visium. Bottom: Expression of NeuroD1 from Allen Mouse Brain Atlas. E17 (https://developingmouse.brain-map.org/experiment/show/100091691), P1 (https://developingmouse.brain-map.org/experiment/show/100091985), P28 (https://developingmouse.brain-map.org/experiment/show/100093575).

Fig. S6. Annotation of Visium clusters in P0 and Adult

Anatomical names of the regions in each cluster were assigned. The Table also contains the names of known marker genes characterizing each cluster.

Fig. S7. Heatmap of the genes specifically expressed in the adult specific clusters

Genes specifically expressed in clusters 6, 7, 8, 10, and 13 in adult brain are shown. See details in Fig. 2.

Fig. S8. NR4A2-positive neurons express independently with Tomato-positive GABAergic neurons in the claustrum

Top: wide-view image of *Dlx5/6*^*Cre;Ai14*^ cortex immunostained with NR4A2. Red cells indicate GABAergic neurons and green cells indicate Nr4a2-positive glutamatergic neurons. Bottom: Expansion of white dot region. Arrows indicate RFP+/GFP-cells.

Fig.S9. Expression patterns of non-VP regions of embryonic thalamus marker genes at P0 (left) and adult (right) stages. Top: Expression of *Gbx2*. Middle: Expression of Tcf7l2. Bottom: Expression of Hsd11b2

Fig.S10. Temporal expression pattern of the embryonic piriform marker gene, Itpka.

