## Supplementary figures and images for "Spatial transcriptome of developmental mouse brain reveals temporal dynamics of gene expressions and heterogeneity of the claustrum"

### Supplementary Figure1

Supplementary Figure 1

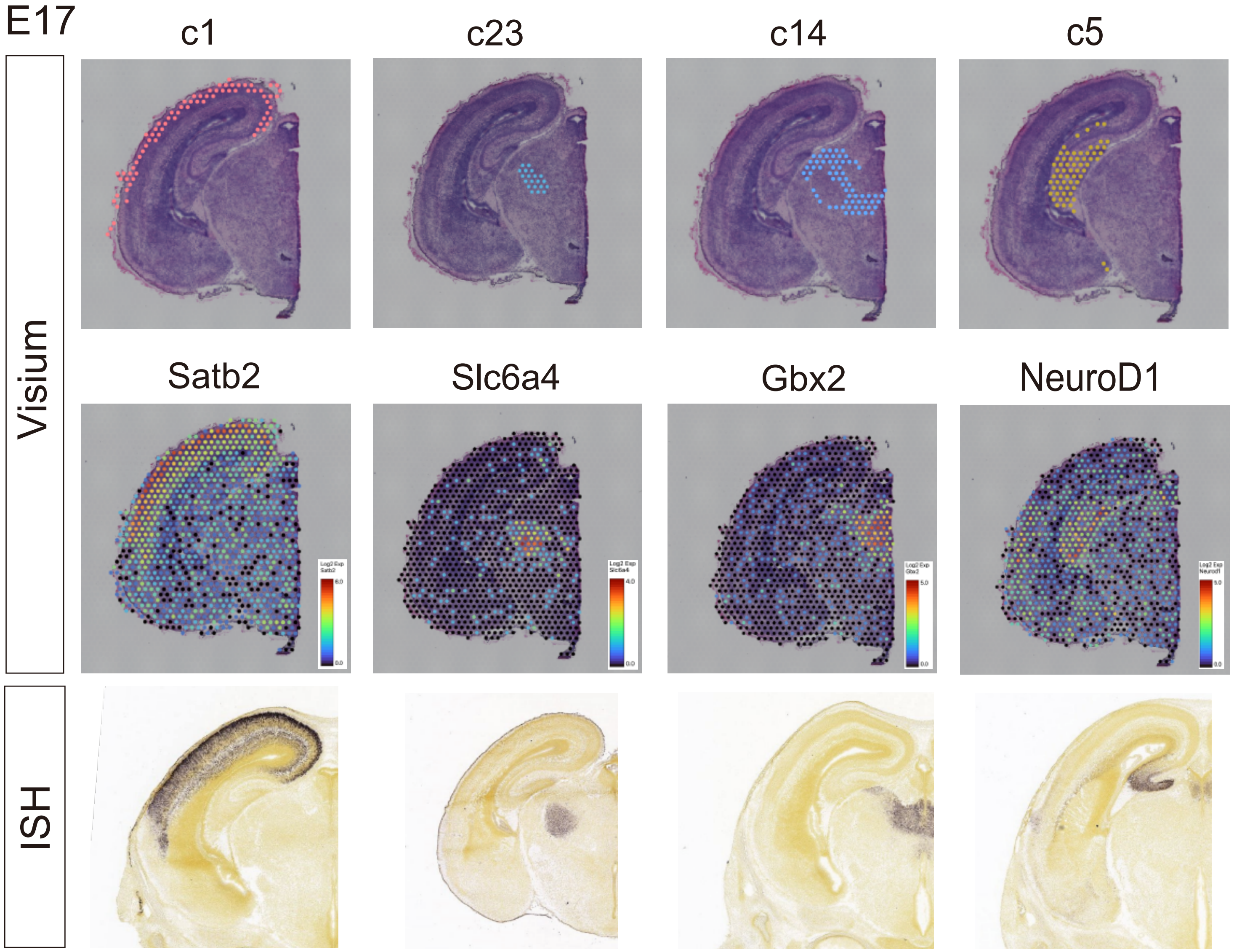

### Supplementary Figure2

Supplementary Figure 2

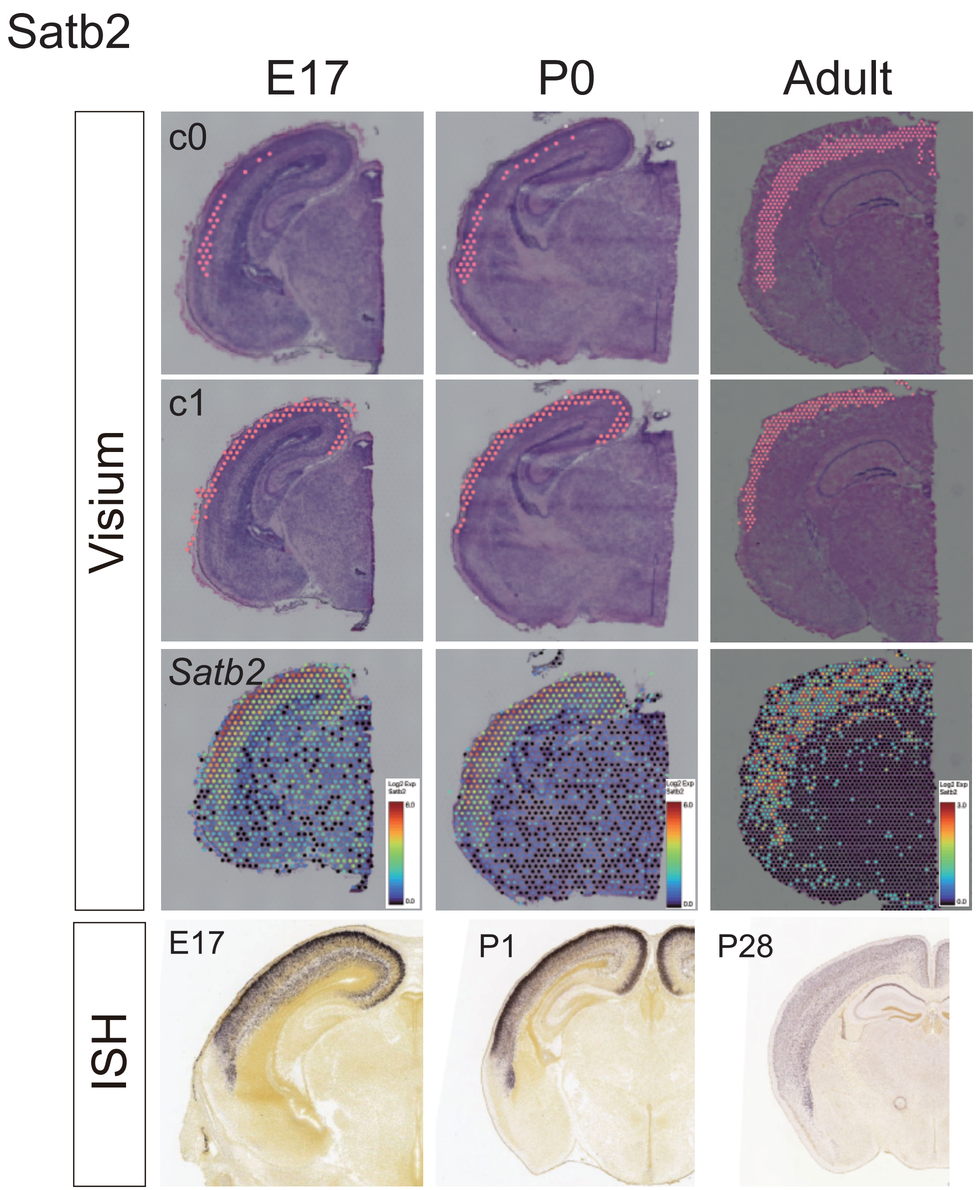

### Supplementary Figure3

Supplementary Figure 3

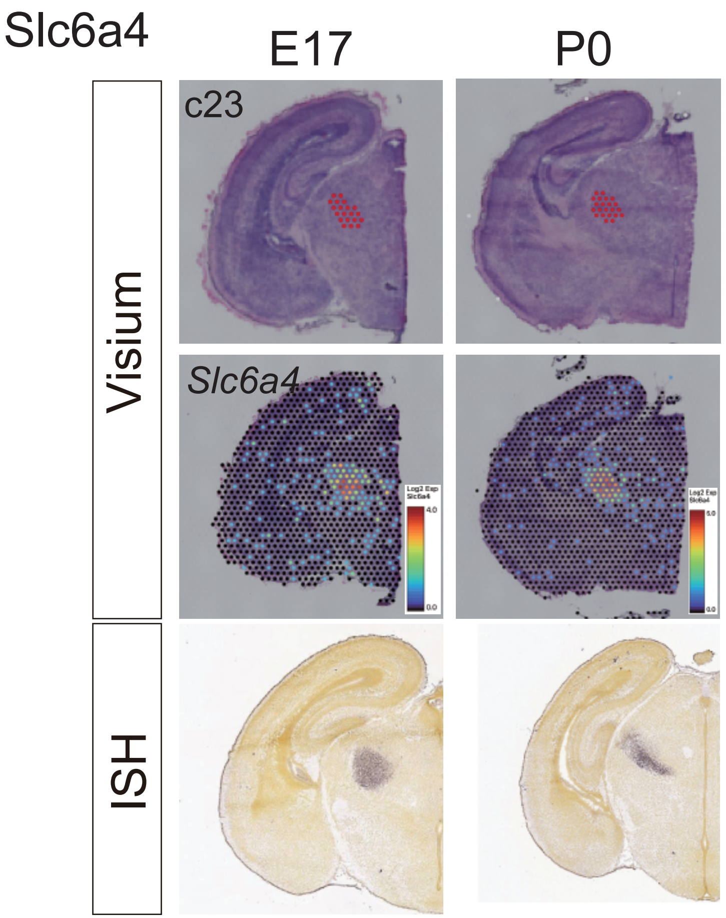

### Supplementary Figure4

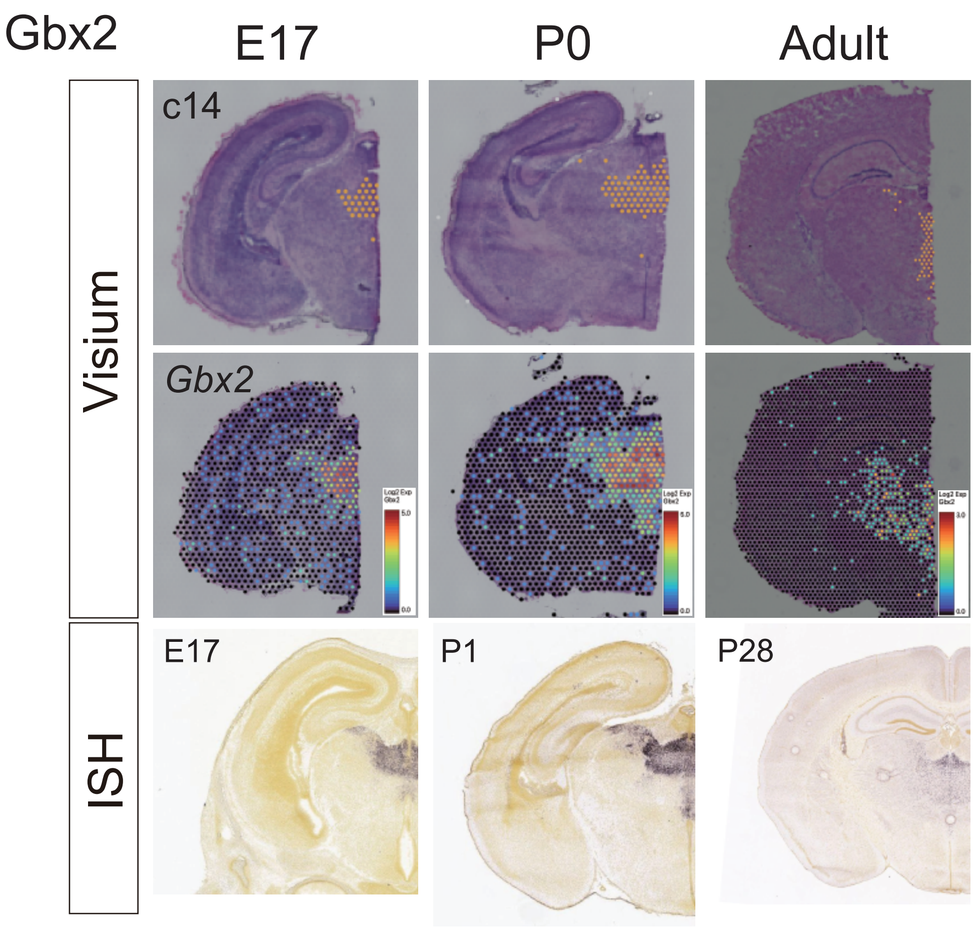

### Supplementary Figure5

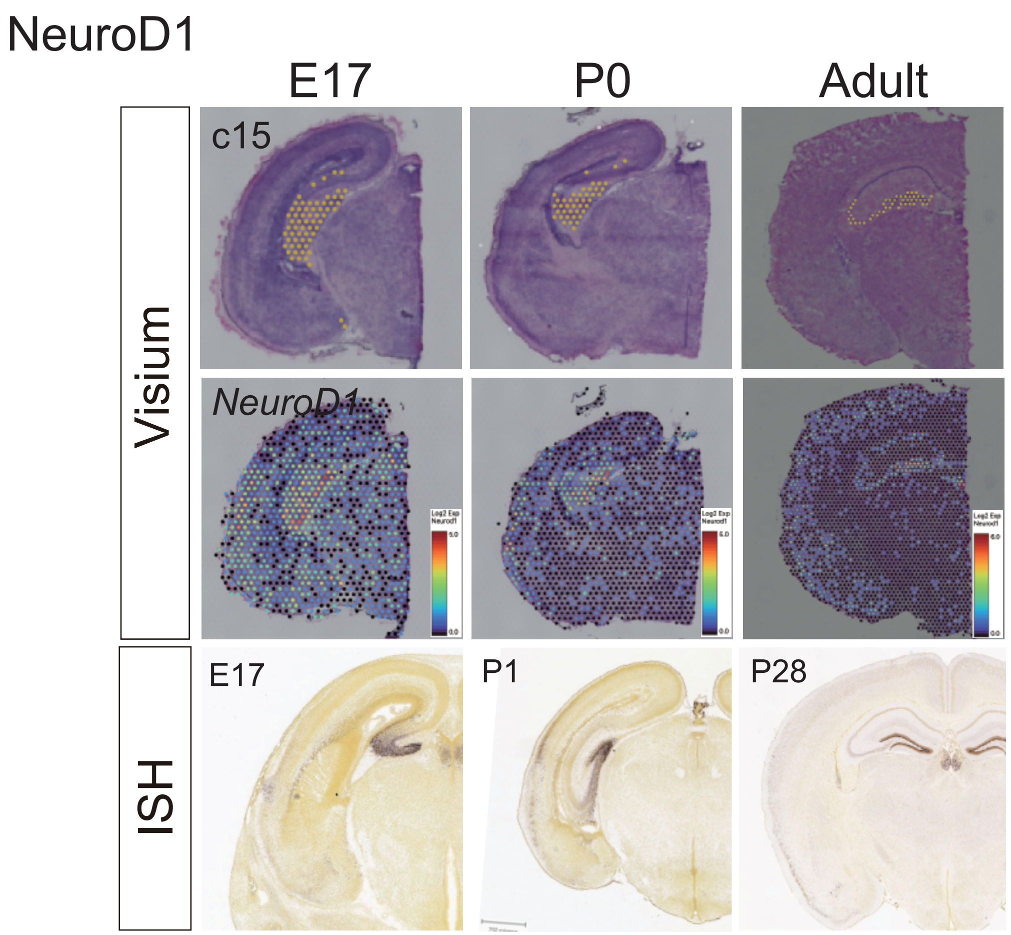

### Supplementary Figure7

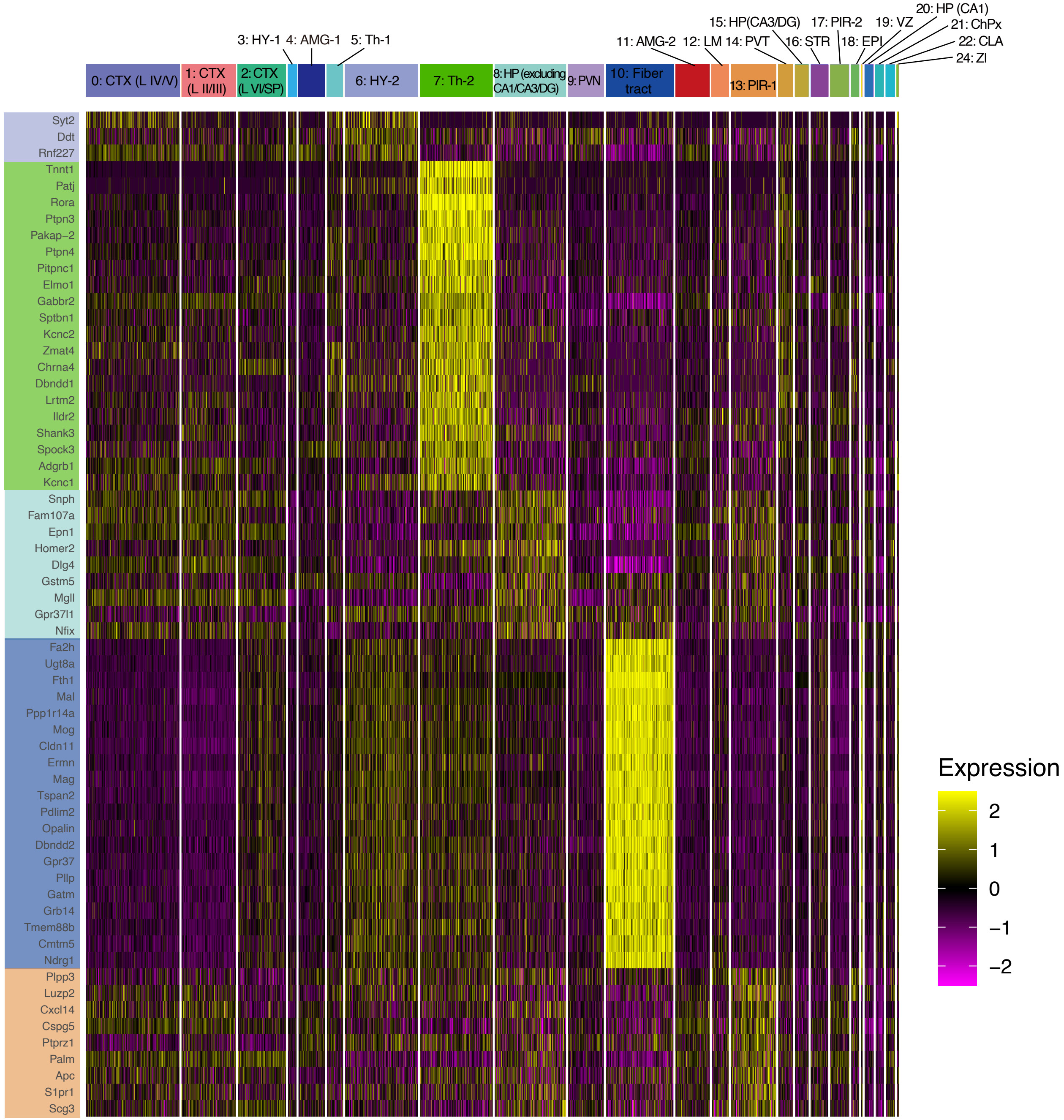

### Supplementary Figure8

Sup_DLT-Nr4a2

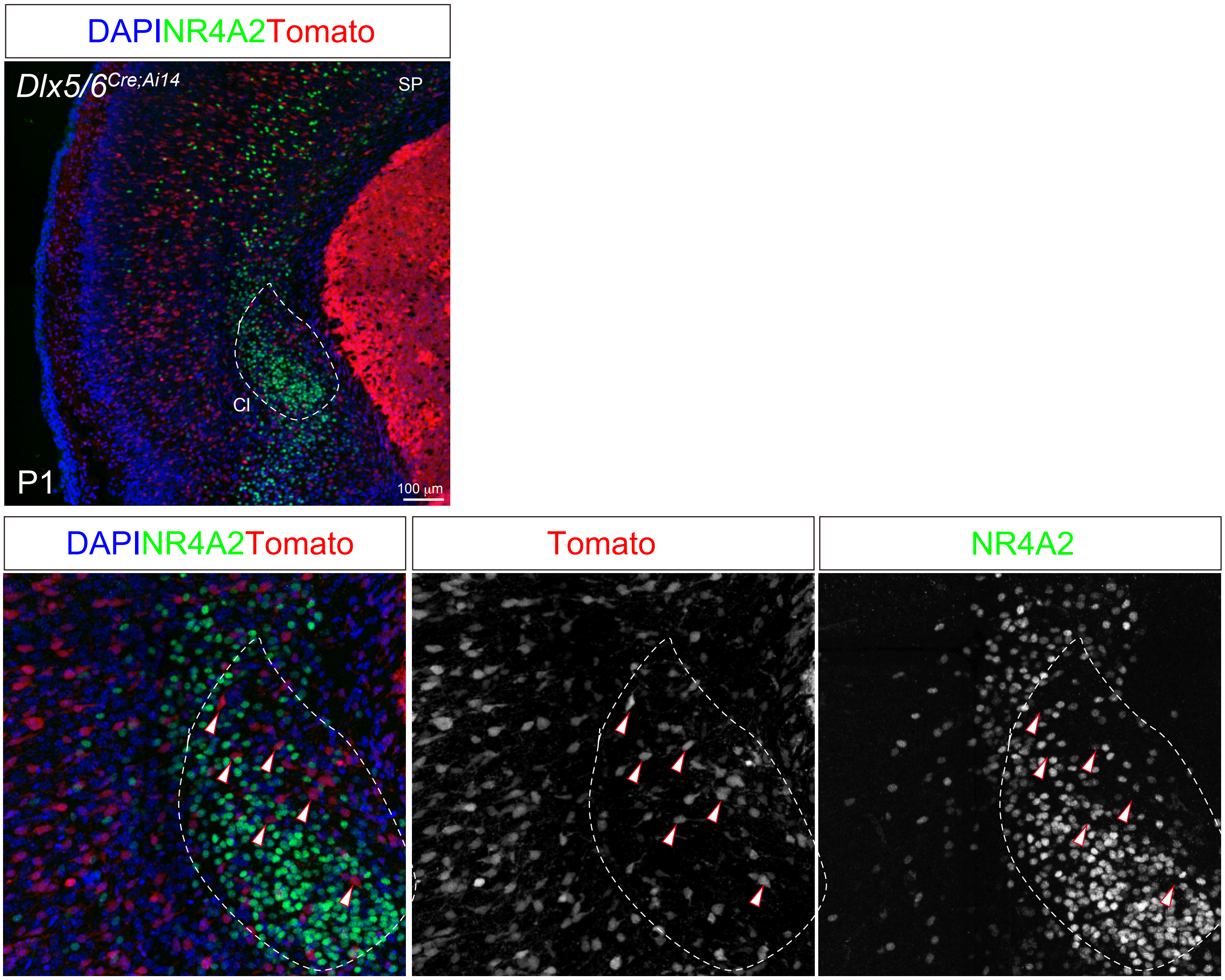

### Supplementary Figure9

Supplementary Figure 9

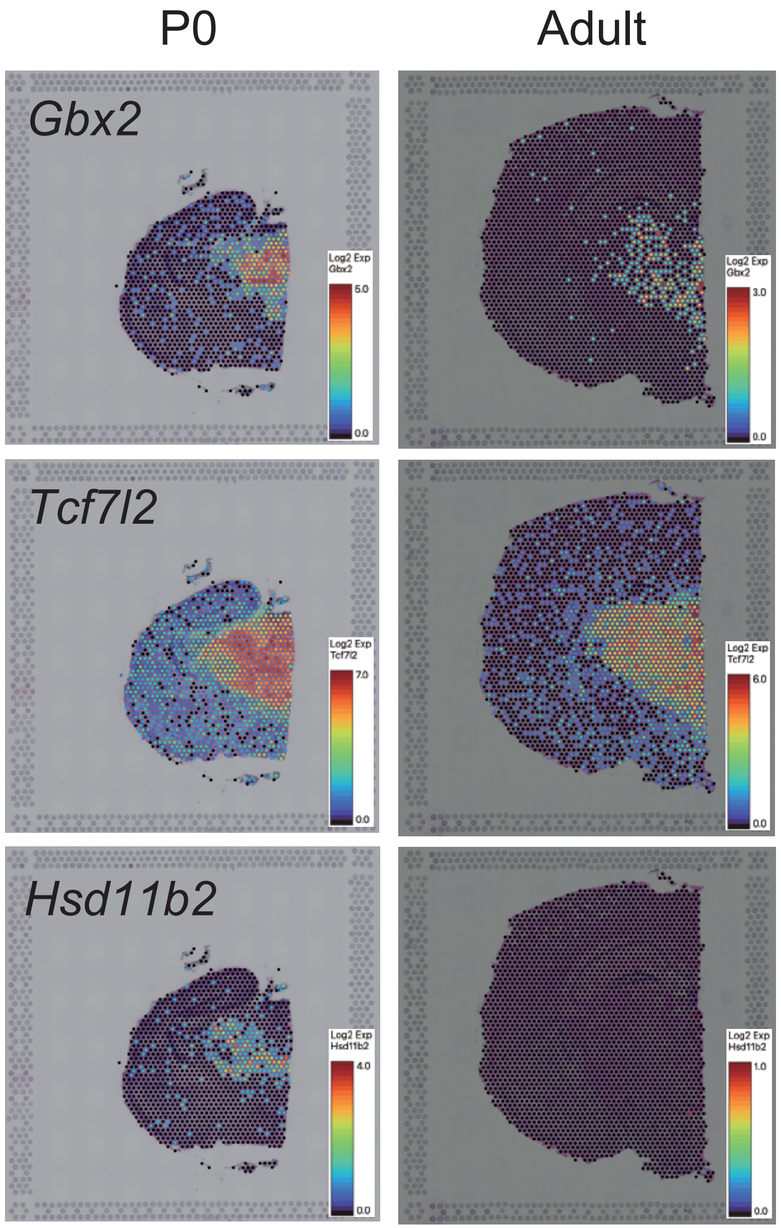

### Supplementary Figure10

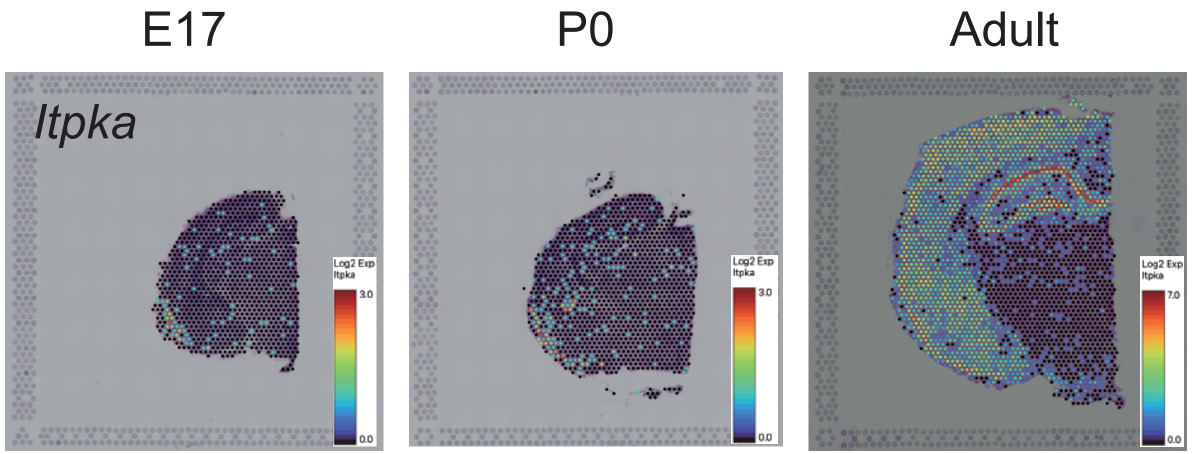
