## Supplementary Figure6 for "Spatial transcriptome of developmental mouse brain reveals temporal dynamics of gene expressions and heterogeneity of the claustrum"

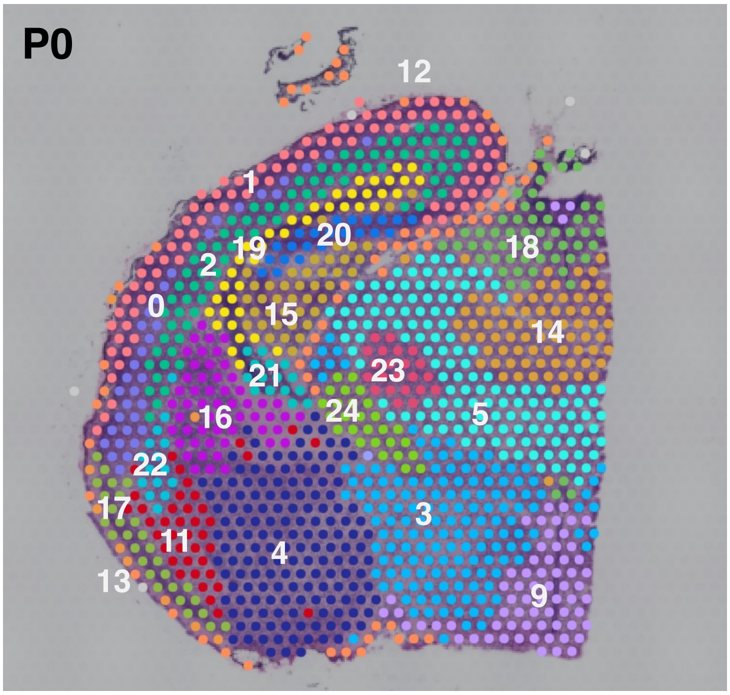


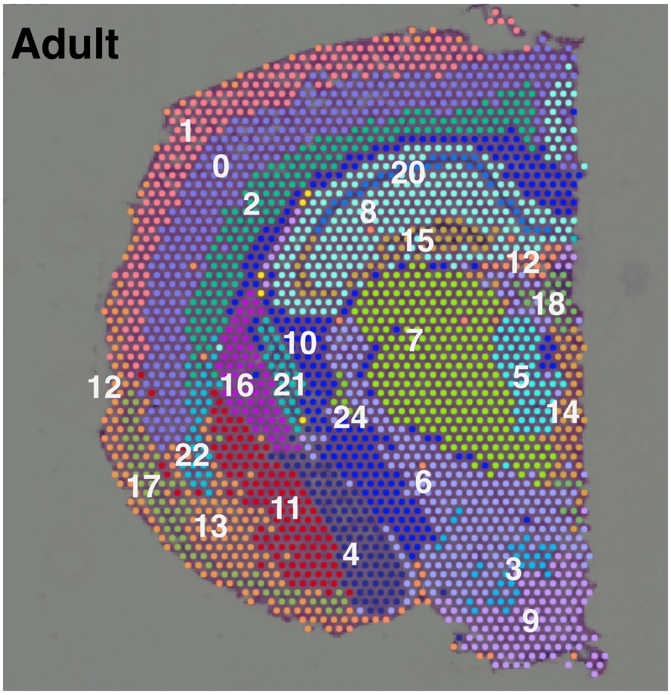


| Cluster# | Annotation | Description | Marker genes |
|---|---|---|---|
| 0 | CTX (L IV/V) | Cortex (Layer IV/V) | Rorb, ETV1 |
| 1 | CTX (L II/III) | Cortex (Layer II/III) | Cux2 |
| 2 | CTX (L VI/SP) | Cortex (LayerVI/Subplate) | Tbr1, Nxph3, Ccn2 |
| 3 | HY-1 | Hypothalamus-1 | Nkx2-2,Otp |
| 4 | AMG-1 | Amygdala-1(medial and central amygdala) | Scn5a, Rasal1 |
| 5 | Th-1 | Thalamus -1 (excluding VP, PVT) | Tcf7l2 |
| 6 | HY-2 | Hypothalamus-2 | Hcrt |
| 7 | Th-2 | Thalamus -2 (excluding PVT) | PRKCd |
| 8 | HP (excluding CA1/CA3/DG) | Hippocampus (excluding CA1/CA3/DG) | Cabp7 |
| 9 | PVN | Paraventricular nucleus | Six6 |
| 10 | Fiber tract | Fiber tract | Plp1 |
| 11 | AMG-2 | Amygdala-2 (lateral, basolateral, basomedial amygdala) | Lypd1 |
| 12 | LM | Leptomeninges | Slc6a13 |
| 13 | PIR-1 | Piriform cortex | Trim54 |
| 14 | PVT | Paraventricular thalamic nucleus | Gbx2 |
| 15 | HP (CA3/DG) | Hippocampus (CA3/Dentate gyrus) | Zbtb20 |
| 16 | STR | Striatum | Adora2a, Ikzf1 |
| 17 | PIR-2 | Piriform cortex | Plxnd1 |
| 18 | EPI | Epithalamus | Pou4f1 |
| 19 | VZ | Ventricular zone | Sox21, Mki67 |
| 20 | HP (CA1) | Hippocanpus (CA1) | Sertm1 |
| 21 | ChPx | Cholpid plexus | Ecrg4, Ttr |
| 22 | CLA/DEn | Claustrum/Dorsal endopiriform nucleus | Nr4a2 |
| 23 | VP (embryo) | Ventral posterior complex of the thalamus (embryo) | Slc6a4 |
| 24 | ZI | Zona incerta | Pvalb |
